# Real-time detection of neural oscillation bursts allows behaviourally relevant neurofeedback

**DOI:** 10.1101/671602

**Authors:** Golan Karvat, Artur Schneider, Mansour Alyahyaey, Florian Steenbergen, Ilka Diester

**Affiliations:** Optophysiology - Optogenetics and Neurophysiology, Albert-Ludwigs-University, Freiburg, Germany; Bernstein Center for Computational Neuroscience, Albert-Ludwigs-University, Freiburg, Germany; Brain Links Brain Tools, Albert-Ludwigs-University, Freiburg, Germany; Faculty of Biology III, Albert-Ludwigs-University, Freiburg, Germany

## Abstract

Neural oscillations are increasingly interpreted as transient bursts, yet a method to measure these short-lived events in real-time is missing. Here we present a real-time data analysis system, capable to detect short and narrowband bursts, and demonstrate its usefulness for volitional increase of beta-band burst-rate in rats. This neurofeedback-training induced changes in overall oscillatory power, and bursts could be decoded from the movement of the rats, thus enabling future investigation of the role of oscillatory bursts.

## Main

Neural oscillations are a frequently reported indicator of neural activity measured invasively via extracellular recordings as local field potentials (LFP), or non-invasively by magnetoencephalogram (MEG) or electroencephalogram (EEG)^1^. In recent years, neural oscillations are increasingly interpreted as transient bursts rather than sustained oscillations^2–5^, and bursts were even suggested as the primary ingredient of all band specific activity^6^. These transient events appear in physiologically relevant time-windows^1^, which makes them optimal candidates to shape behaviour in a trial-by-trial fashion^7^. Despite the increasing attention to these transient bursts, their role in neural computation, and ultimately in producing behavioural outputs, remains controversial^3^.

If indeed bursts of neural oscillations play a role in behaviour, we hypothesize that 1. subjects can learn to increase the burst rate in a goal-directed manner (i.e., neurofeedback, Fig. 1a), 2. burst-rate-increase will lead to global (averaged) power increase, and 3. burst occurrences can be predicted based on behavioural readouts.

**Figure 1.**
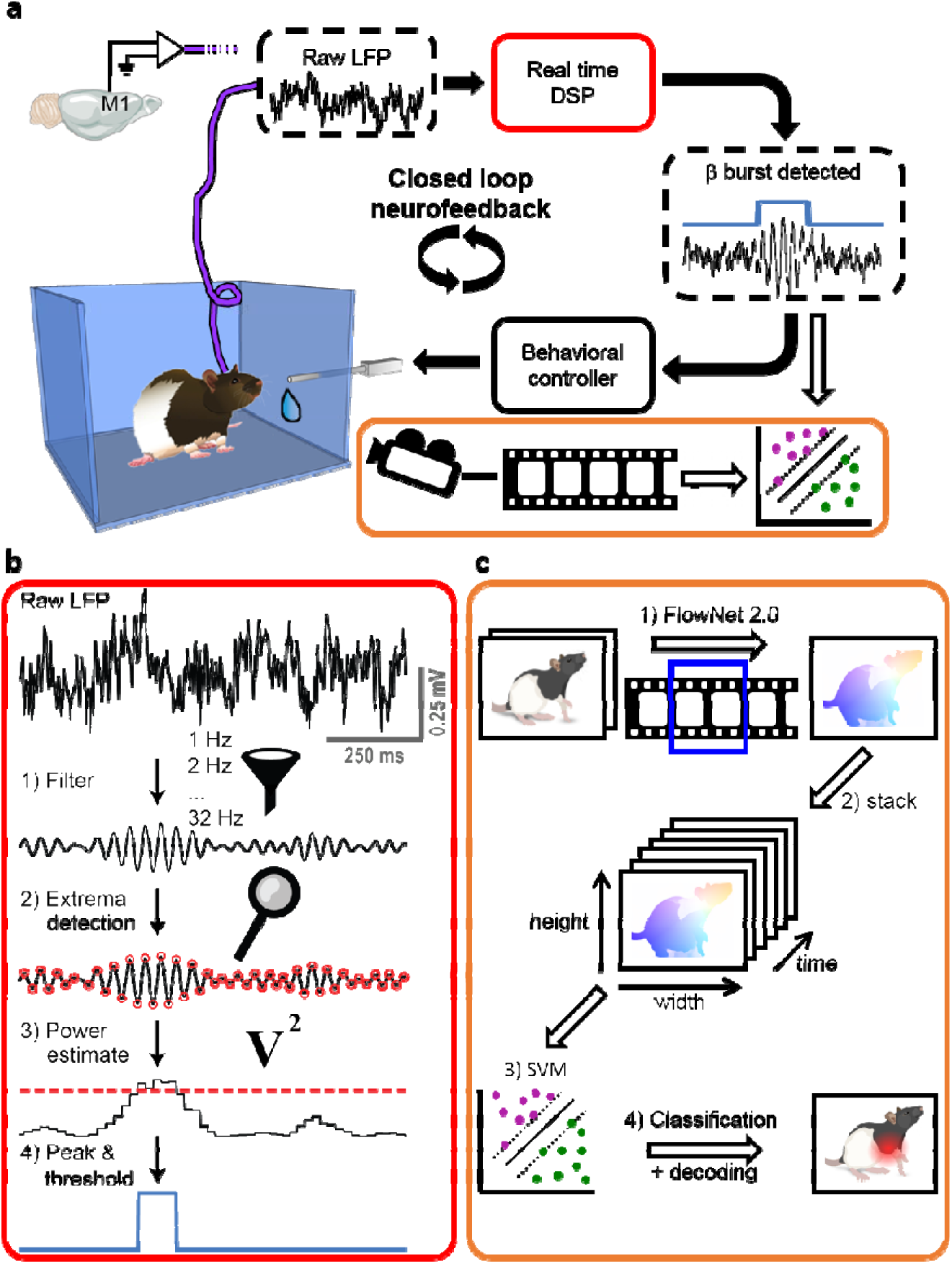
Overview of LFP β-event based neurofeedback method. **a**. The Setup. LFP signal from the motor cortex of a freely moving rat was measured and fed into the real-time digital signal processing unit (DSP, red outline). Upon detection of an LFP beta-burst, the rat was rewarded with sucrose water. The activity of the rat was videotaped in synchronization with the electrophysiological data, and videos were analysed offline by a machine-learning algorithm to detect movements indicative of beta bursts (orange outline). Black arrows: online analysis. White arrows: offline analysis. **b**. Real-time LFP-burst detection algorithm. 1) The raw signal was filtered by an array of digital narrowband finite impulse response filters. 2) Extrema points were detected in the filtered signal. 3) The square of the amplitude in an extrema point was latched until the detection of the next extrema point and served as an estimate of power. 4) If the power in a specific frequency was higher than the power of the frequency above and the frequency below, as well as the value of the 98^th^ percentile of the target frequency power calculated online, it was defined as a burst. A burst was rewarded if it happened in the targeted frequencies, and lasted ≥70 ms. **c**. Offline algorithm for decoding behaviour. A support vector machine (SVM) model was trained to classify epochs with or without beta bursts (as detected by the real-time DSP). Movements of the rat (1) were approximated via optical flow, calculated from adjacent frames with FlowNet 2.0^18^. A time stack of flow-images (2) was used as an input for the SVM-classifier (3). Classification accuracy and attention (distance to the decision function) were used to evaluate the model in the temporal and spatial domains (4).

As a test-bed for these hypotheses, we chose the rat motor cortex and focused on LFP bursts of beta-waves (15-30 Hz), which have been documented in humans, monkeys, and mice. These short bursts (<150 ms) have been correlated with memory, movement and perception^4–6,8,9^, and reported to be correlated with behaviour up to 300 ms relative to the burst occurrence^8^.

However, a method to measure these short-lived bursts in real-time for addressing these hypotheses is missing. The first challenge in developing such a method is formally defining LFP bursts. We suggest defining a burst as a power peak in time and frequency, exceeding a threshold^5^. When defining the threshold, two key points have to be addressed: first, it should be calculated from the ongoing session to represent the current brain-state of the subject, as the global LFP-power can change between subjects and sessions. Second, it should be based on a defined percentile, as opposed to central tendency measures (i.e., mean and median), to assure a statistically sound significance definition under non-normal distributions.

The second challenge is the detection of such short-lived peaks, which requires minimal pre-processing and delay, as well as high time and frequency resolutions. Here we present a real-time digital signal processing (DSP) method, capable to detect short and narrowband bursts (Fig. 1b). The algorithm is based on 32 real-time narrow bandpass filters. These digital finite-impulse-response (FIR) filters are centered on steps of 1 Hz. In real-time, the acquisition system detects peaks and troughs in the filtered data, and determines the power based on the amplitude of these extrema. As both peaks and troughs are taken into account, the time resolution is half the period of each frequency. The FIR filters ensure that there is no distortion due to the time delay of frequencies relative to one another (i.e. linear phase), resulting in a fixed delay of 130 ms for each frequency (see movie S1). This allows a direct comparison of neighboring frequencies necessary for peak detection, at unprecedented frequency and temporal resolutions (1 Hz and 130ms + half the period of each frequency), outperforming conventional online methods (Fig. S1). To close the loop between oscillatory events and behaviour, we linked the DSP system with an operant conditioning apparatus for rodents, and synchronized videos with the LFP-recordings for offline behavioural analysis (Fig. 1c).

For demonstrating the efficacy of the real-time method and investigating whether rats can volitionally increase the rate of beta-bursts, we implanted laminar probes in the motor cortex of 3 rats. The freely moving rats were placed in the closed-loop neurofeedback apparatus, where artefact-free LFP was measured and analysed in real-time. Occurrences of oscillatory bursts in one of the frequencies in the beta band (20-25 Hz for 2 rats, 15-20 Hz for 1 rat), higher than the 98^th^ percentile of power and longer than 70 ms were rewarded (movie S1).

Within 9 sessions of training, oscillatory bursts became identifiable in raw LFP traces (Fig. 2a and 2b), accompanied by a 34% increase in general (averaged) beta power (Fig. 2c, p=1.25*10^−12^, ANOVA). The method allows to follow the learning of the rats in a session-by-session manner. Both power (Fig. S2a) and number of rewards (Fig. S2b) were linearly correlated with session progress (ρ=0.52, p=0.0059 for power and ρ=0.49, p=0.0101 for rewards). Each rat had one prominent session with a sudden power increase (“aha-effect”, Fig S3, p<5.19*10^−6^, ANOVA with Bonferroni correction). This power increase occurred in each of the targeted frequencies in the beta range with 1 Hz resolution (Fig. S4). Importantly, the number of rewards after the power increase was 22% higher than beforehand (p=2.14*10^−4^, two-sided t-test, Fig. 2d), indicating that rats volitionally increased the beta-bursts-rate and -power by neurofeedback training. The average of beta-power across the full 30 minutes of each session and the number of rewarded short-living-bursts were highly correlated (ρ=0.89, p=5.9*10^−10^, Fig. 2e). These findings strongly support the critical influence of bursts on the global (averaged) beta-band power, as was suggested previously^6^.

**Figure 2.**
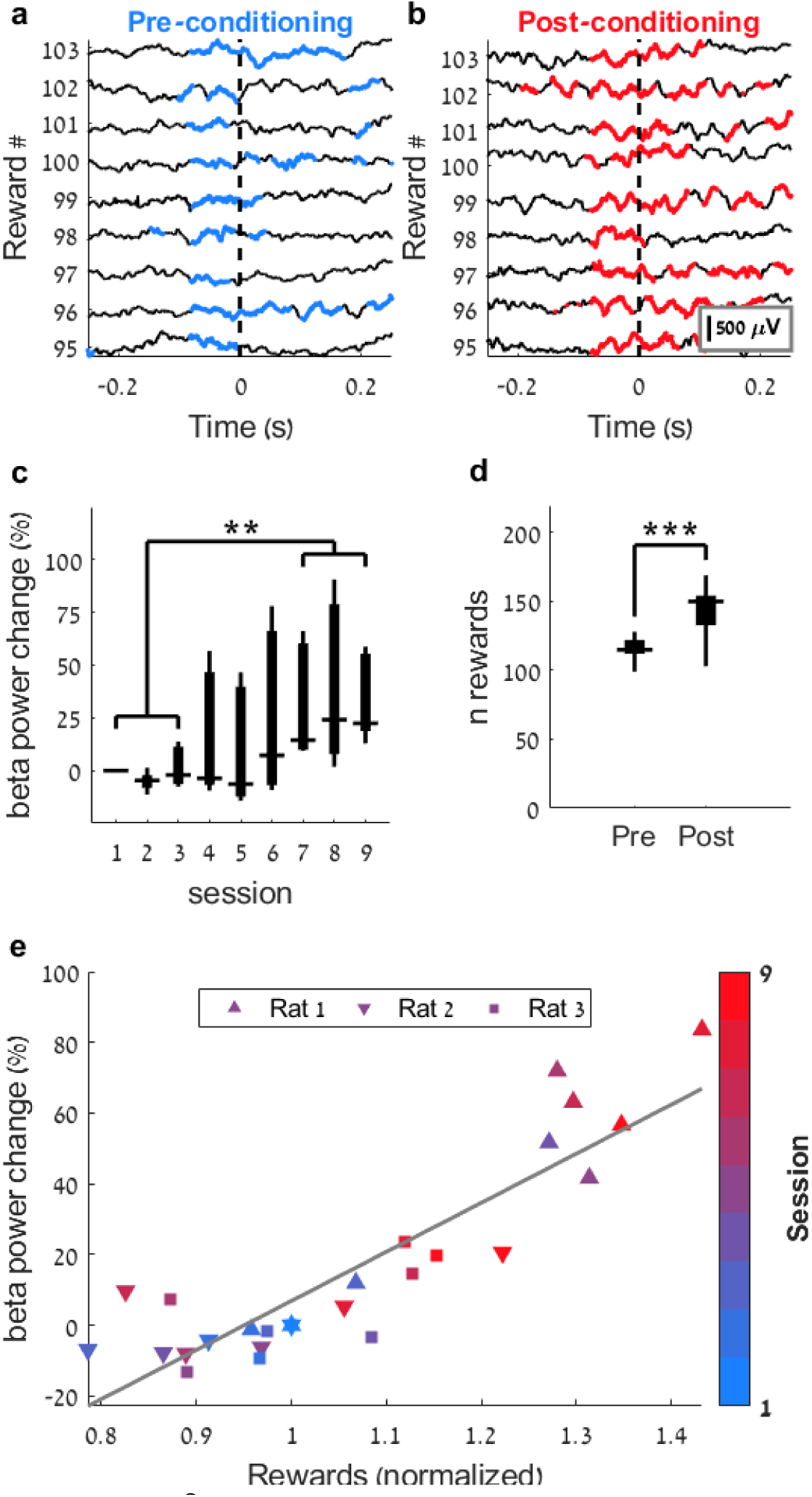
Neurofeedback increases β-burst power and rate. Raw LFP traces before (**a**) and after (**b**) 9 sessions of neurofeedback training reveal that the rat was conditioned to exhibit oscillations. Time-points in which beta-power exceeded the 98^th^ percentile threshold are marked in blue (**a**) or red (**b**). Reward was delivered at time = 0. **c**. Power change in the targeted beta frequencies relative to the power on day 1. Two-way ANOVA (factors: session and frequency), effect for session: *F*_(8,10)_=9.33, p=2.65*10^−10^. Bonferroni corrected post-hoc analysis shows significant difference between sessions 1-3 and sessions 7-9. **-p<0.01. **d**. Number of rewarded beta-bursts before (pre) and after (post) the identifiable session of power increase (“aha-moment”, see details in figure S3). ***-Two-sided t-test *t*_(25)_=4.32, p=2.14*10^−4^. In **c** and **d**, the presented elements are: centre line, median; box limits, upper and lower quartiles; whiskers, full distribution. **e**. Correlation between number of rewards relative to day 1 and beta power change as in **c** for each rat in each session. Colours indicate the session number and each rat is denoted with a different marker. Pearson’s ρ= 0.89, p=5.9*10^−10^.

In order to test for a link between the detected LFP bursts and behaviour, we analysed movements as behavioural readout, since we recorded from the motor cortex. Therefore, we performed video recordings of the behaviour of the rats in synchronization with the LFP recordings. A critical matter for behavioural analysis is to avoid bias and maintain time-scale accuracy relevant to the underlying brain activity (tens to hundreds of milliseconds for LFP-bursts^1,8^). For a human observer, it is almost impossible to fulfil these criteria. Recently, it was suggested that application of machine learning analysis approaches can overcome these difficulties^10^. Therefore, we trained a support vector machine (SVM) supervised learning algorithm to decode the occurrence of neuronal LFP bursts from the videos in an offline manner (see Fig. 1c). We were able to link beta-bursts to behaviour, as the SVM supervised-learning algorithm could reliably decode occurrences of bursts based on the rats’ movements with an 18% better prediction accuracy for true positive epochs compared to shuffled (Welch t-test, p=0.03, Fig. 3a). The unbiased attention of the network (see methods) increased during the trial towards burst initiation (ρ=0.87, Fig. 3b), supporting the current view of increased beta oscillations power at the termination of movements^11^. Additionally, the attention in space was set to the frontal body parts of the rats (e.g. snout, Fig. 3c, right panel), indicating that indeed the rats’ movements were important for decoding LFP-bursts from the videos.

**Figure 3.**
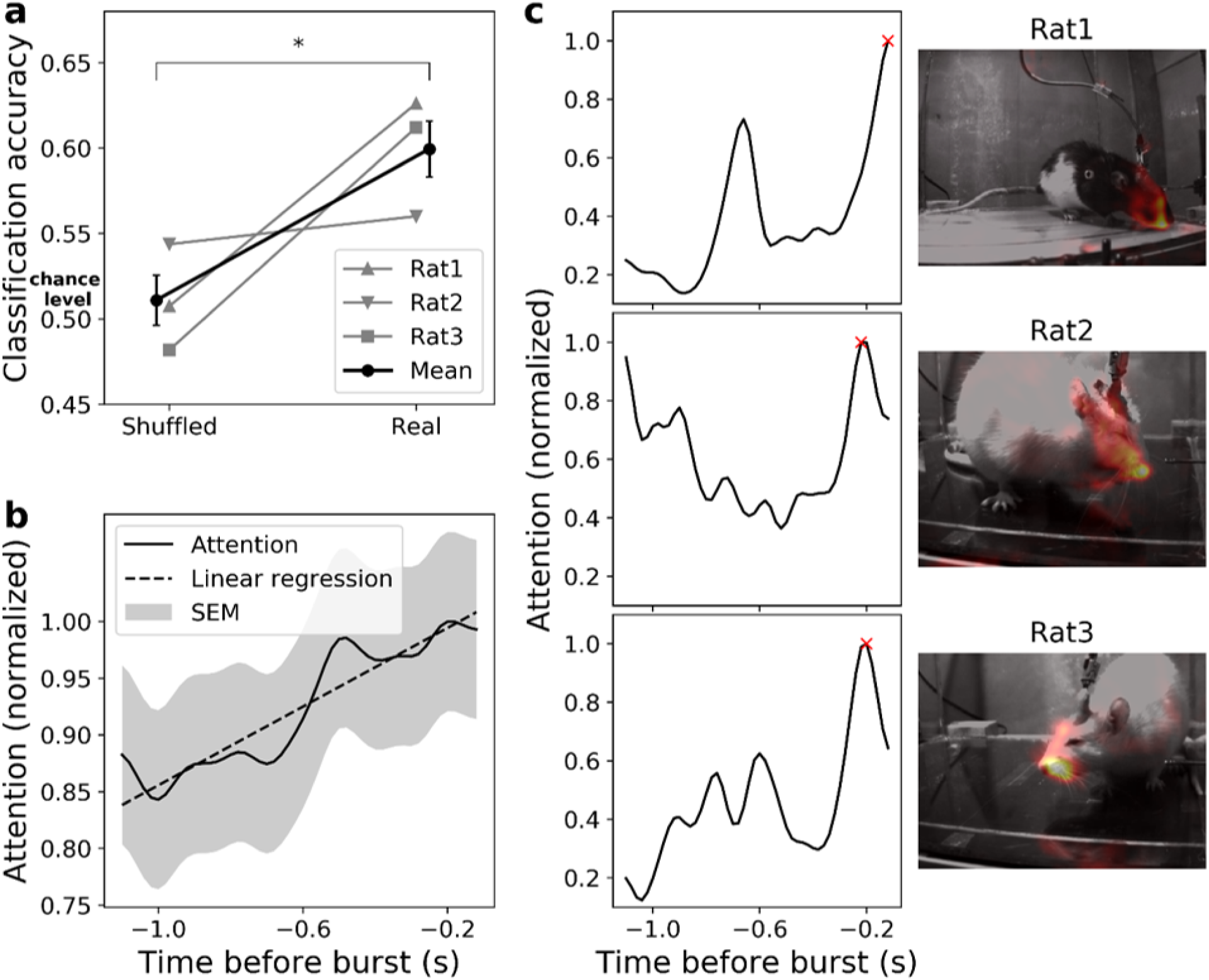
Behavioural effects of β-event neurofeedback training. **a**. Classification accuracy of the SVM models (n=3 repetitions) trained on epochs with correct identity vs. models trained on epochs with shuffled identity. SVM models achieved above chance accuracy (18% increase, 0.6 vs 0.51, two-tail Welch t-test, t_(4)_=3.284 p=0.03). **b**. Temporal attention of the models on correctly predicted true samples. The time course of the mean normalized attention over all true positive epochs ± SEM indicates that movements leading to beta bursts occurred primarily shortly before burst initiation (Linear regression, ρ=0.87, p=2.8E-23). **c**. Examples of the model attention in individual epochs. The temporal (left panel) and spatial (right panel, heat-map overlaid on the video frame with highest attention, marked with red x in temporal profile) attentions of the model follow similar patterns for representative epochs of each rat. The attention implies that the movements of frontal body parts, shortly before burst initiation, were used to predict the epoch class.

Here, we introduce a real-time LFP-burst-based neurofeedback system in freely moving rodents. Previous animal studies have employed spike detection^12^, Calcium transients^13,14^, and sustained LFP oscillations for neurofeedback^15,16^. Our results demonstrate for the first time the potency of real-time LFP transient burst detection for neurofeedback. Furthermore, we confirm the impact of bursts on global oscillatory power and behaviour. We focused on detecting and manipulating beta-bursts in the motor cortex, but the algorithm is flexible and could be adjusted to target bursts in other frequency ranges and brain areas. Thus, our approach can be a starting point for a plethora of studies targeted at understanding the causal role of oscillatory bursts. For example, instead of artificial external stimuli, real-time burst-triggered stimulus presentations could be combined with behavioural and electrophysiological measurements, thereby allowing to probe the intrinsic function of oscillatory bursts. Furthermore, neurofeedback has been clinically used for decades without a clear understanding of the underlying neural mechanisms^17^. As our tool is ideally suited for rodents, it can be combined with additional invasive or non-invasive treatments and post-mortem histology, thereby providing a new testbed with high relevance for future clinical developments, e.g., to advance the design and patient training of brain-machine-interface prosthetic devices^17^.

## Supporting information

Video of a freely moving rat with raw LFP trace and power estimation

## Acknowledgements

This work was supported by the Deutsche Forschungsgemeinschaft (DFG) through the Clusters of Excellence BIOSS BrainLinks-Brain-Tools (EXC 1086) to I.D. and the ERC Starting grant OptoMotorPath (338041) to I.D. We thank Michael Tangermann for discussions and advice for the design of the real-time feedback algorithm, David Eriksson and Philippe Coulon for contributions to establishing the electrophysiological setup and helpful remarks on the manuscript, Thomas Brox for providing the FlowNet algorithm and feedback on previous versions of the manuscripts, Patrick Ruther for providing laminar probes, Izhar Bar-Gad and Ileana Hanganu-Opatz for advice on artefact avoidance in LFP recordings, and Tucker-Davis-Technologies for the assistance in designing the real-time circuit.

## Contributions

G.K. conducted surgeries, animal training, electrophysiological recordings and data analysis, built the setup, and designed the algorithm and circuit. A.S. and F.S. conducted the machine learning analyses. M.A. conducted surgeries. G.K. and I.D. designed the study and wrote the manuscript.

## Movie S1: Video of a freely moving rat with raw LFP trace and power estimation (related to figure 1)

Example video of the freely moving rat (left) with a raw LFP trace (bottom right) and the power estimation as computed online (spectrogram, top right). The colour map is normalized to the 98^th^ percentile of the power in each frequency, i.e., values higher than 1 are above the statistically defined threshold. If the power in a specific frequency crossed the threshold and was also higher than the neighbouring frequencies, it was considered a burst and denoted with a white overlay (over the spectrogram) or red overlay (over the LFP trace). If a burst lasted >70 ms, the rat was rewarded with sucrose water (blue lines above the LFP trace). The next burst could be rewarded only after the end of reward delivery. Note that the delay between the LFP trace and the power estimation is constant at 130 ms, which is due to the group delay of the online filters.

## Methods

### Animals and surgery

In this study, we used adult female rats (n = 3, 56 ± 5 weeks of age, 351 ± 21 g, mean ± standard deviation at surgery day, two Sprague Dawley and one Long Evans, Charles-River, Sulzfeld, Germany, table S1), which were housed under an inversed 12 h light dark cycle. We implanted 32 IrOx electrode silicone probes (1 shaft, 50 μm between electrodes, gift from Patrick Ruther, Technical Faculty, University of Freiburg) in the left motor cortices (2.4 mm lateral and 1.5 mm anterior to bregma). To anaesthetize the rats, we injected 80 mg/kg Ketamine (Medistar, Holzwickede, Germany) and 100 μg/kg Medetomidine (Orion Pharma, Espoo, Finland) intraperitoneally, as well as 10 mg/kg Carprofen (Rimadyl, Zoetis, Berlin, Germany) and 25 μg/kg Buprenorphine (Selectavet, Dr. Otto Fischer GmbH, Weyarn/Holzolling, Germany) as analgesics. To maintain vital body measures, a heating pad connected to a rectal temperature sensor (Stoelting, Dublin, Ireland) maintained the rat’s body temperature at 37°C, and a pulse oximeter (model 2500A VET, Nonin Medical, Plymouth, MN) monitored the blood oxygen level and heart-rate while delivering oxygen-enriched air (1 l/min) through a face mask. After placing the rat in the stereotactic frame (David Kopf Instruments, Tujunga, CA) and exposing and cleaning the skull, we thinned the bone above motor cortex with a dental drill (MH-170, Foredom, Bethel, CT). A final small (∼1mm) craniotomy was made over a cortical area with no large blood vessels. We connected the flexible wire ribbon of the probe to an adaptor compatible with Tucker-Davis-Technologies (TDT, Alachua, FL) headstage’s zero-insertion-force (ZIF) connector, and held the ribbon on the stereotactic frame by a vacuum holder (Atlas Neuroengineering, Leuven, Belgium). As reference and ground, we connected 130 μm diameter silver wires (Science Products, Hofheim, Germany) and wrapped them around self-tapping screws (J.I. Morris Company, Southbridge, MA) positioned above the cerebellum. After lowering the probe until the tip reached 2 mm below dura, we applied a Kwik-Cast Sealant (World Precision Instruments, Sarasota, FL) over the craniotomy and a thin layer of super bond C&B cement (Sun Medical, Shiga, Japan) over the implant and supporting skull-screws. Afterwards, we added several layers of Paladur dental cement (Heraeus, Hanau, Germany) to cover the probe and adaptor, leaving only the ZIF connector of the adaptor exposed. To protect the connector, we attached a metal 780-11 paper-clip (ALCO, Arnsberg, Germany) to the adaptor. After the surgery, we placed the rat in a heated, oxygen-enriched chamber until it woke up, and administered Carprofen (10 mg/kg) and Buprenorphine (25 μg/kg) daily for 3 days. Rats were given >7 days to recover from surgery before water-restricted training began. All procedures were in accordance with the guideline RL 2010 63 EU and were approved by the Regierungspräsidium Freiburg.

### Real time burst detection algorithm

#### Data acquisition and filtering (Fig. 1B)

Real-time analysis of signals as small as a few microvolts demands artefact-free recordings (see table S2 for artefact sources in electrophysiological recordings from freely moving animals and the measures taken to reduce their influence to a minimum). We acquired raw broadband signals at 25 kHz using a digital headstage (ZD32, TDT) and down-sampled them to 1 kHz. One electrode, located at a depth of 1100 μm, was selected for analysis. Filtering of the raw signal took place within the online digital signal processor (RZ2 BioAmp, TDT), by Bartlett window finite impulse response (FIR) filters, with order of 256, stop-band attenuation of 6 dB and pass-band frequency width of 1 Hz. The filters were centred on frequencies ranging from 1 to 32 Hz (in steps of 1 Hz). We generated the filter coefficients with the Matlab (Mathworks, Natick, MA) function “fir1”, as follows: b = fir1(N, [Fc1 Fc2]/(Fs/2), ‘bandpass’, win, flag) with b the array of coefficients and the following parameters: order (N) = 256; Fc = central frequency of interest (1 to 32 Hz); lower limit of pass-band (Fc1) = Fc-0.5 Hz; upper limit of pass-band (Fc2) = Fc+0.5 Hz (resulting in pass-band of 1 Hz); sampling frequency (Fs) = 976.5625 Hz (∼1 kHz as implemented in the TDT system); window (win): Bartlett window of order N+1 generated by the Matlab function “bartlett”; normalization option (flag) = ‘scale’ (the magnitude response of the filter at the centre of the passband is 1). The delay of the filter (group delay + computation time) was 130 ms for each frequency, allowing direct comparisons between frequencies. Two video cameras (Basler acA640-750um) recorded the rats’ movements from orthogonal viewpoints. To ensure that video frames were in synchrony with the electrophysiological data, the acquisition system triggered the cameras via a TTL signal (50 Hz square wave with 40 μs width).

#### Power and phase estimation

The real-time algorithm detected extrema points in each frequency by applying a peak/trough feature detection routine on the filtered signal. The square of the LFP amplitude upon extrema-point detection served as power estimate. Latching the power value until the next extrema point detection yielded a time resolution of half the period time for each frequency. Although phase estimation was not used in the current study, the algorithm estimated the phase by forcing a saw-tooth signal with amplitude 2π and frequency f to 0 upon detection of a peak and to π upon detection of a trough. To align the determined phase to the true phase, the phase-reset was delayed by δ(f)=⌈d/T⌉*T-d, where δ(f) is the delay in seconds to the next peak, f frequency in Hz, d group delay of the filter in seconds, T time period in seconds and ⌈ ⌉ the ceiling operator. This delay compensated for the filter group delay, and aligned the phase to the next assumed peak of the real signal.

#### Artefact rejection

Rare events of high amplitude LFP deflections (which occurred mainly when the rats crunched their teeth or touched the headstage during grooming) have extremely high power in a large range of frequencies. These artefacts might be erroneously interpreted as bursts, and have a tremendous effect on the power distribution. Therefore, LFP values from the selected electrode which exceeded a threshold of 500 μV were considered artefacts, and data points 500 ms before and after the detected artefact were removed from analysis. When an artefact was detected, the rat received 1 second of 90 dB SPL white noise and a no-reward timeout, to train it to avoid causing artefacts. To avoid interpreting high frequency action potentials or very low frequency trends as artefacts, the raw LFP went through a 12 dB per octave Butterworth bandpass filter between 2 and 250 Hz prior to thresholding. Overall, 0.0057±0.001 (mean ± SEM) of samples per session were rejected.

#### Burst detection

The real-time DSP buffered the power in each frequency together with detected artefact times and sent them to a Matlab routine for dynamic calculation of the percentiles every second. The Matlab routine used the last 15 acquired seconds for calculation, while ignoring LFP power values during artefact rejection times. As substitute for values during artefact rejection times, the routine used earlier values, assuring full 15 seconds of artefact-free data for percentile calculation. The power values of the target percentile in each frequency were sent back to the DSP for real-time burst detection; in every time point, the DSP compared the power value in each frequency to the target power percentile as well as to the power of the neighbouring frequencies. If the power in a target frequency range (15-20 Hz for 1 rat, 20-25 Hz for 2 rats) exceeded these values, the DSP sent a TTL pulse to the behavioural controller (Med Associates, Fairfax, VT), signalling a burst.

### Neurofeedback training

To obtain clear video images and to avoid artefacts caused by electrostatic discharges, we built an open-top glass cage sized 30 × 26 × 40 cm (width × length × height) and positioned it inside a grounded Faraday cage. A 2 × 12 mm infusion cannula (1464LL, Acufirm, Dreieich, Germany) served as a spout for 3% sucrose water delivery as reward for the water deprived rats, and was controlled by an infusion syringe pump (PHM-107, Med Associates, see Fig. 1A). In order to allow time for the initial period of percentile computation, in the first 15 seconds of the session, the rats received 5 rewards of 50 μl sucrose water delivered every 3 seconds. Henceforth, each session lasted 30 minutes. Upon detection of a rewarded burst from the DSP longer than 70 ms, the rats obtained 30-75 μl sucrose water rewards delivered at 50 μl/ second. The reward size was adjusted to ensure that the rat received 8-14 ml water per day, and was accompanied by a 12 kHz, 90 dB SPL pure tone to facilitate learning. During reward delivery and 1 second after a reward or an artefact, no reward could be obtained. Training lasted 9 sessions (1-2 sessions per day) during the dark period. We weighed the rats before each session to assure they stayed above 80% of their pre-deprivation weight.

### Offline machine learning and video analysis

#### Flow calculation

To relate the occurrence of beta bursts to behaviour, we analysed the apparent movements of the rats using optical flow (figure 1C). FlowNet 2.0 (https://github.com/lmb-freiburg/flownet2) calculates the pixel changes between two images, resulting in an x- (u) and y- (v) vector components of every pixel between two consecutive images. Individual frames from one of the cameras were extracted via ffmpeg (2.8.15, https://github.com/FFmpeg/FFmpeg), scaled down to 320 × 240 pixels and passed through FlowNet 2.0 to calculate the optical flow between the frames.

#### Data preparation

Time points of beta bursts as detected online were used to extract the corresponding frames. We used 50 frames (corresponding to 1 second) from 1.1 to 0.1 second before the time of the beta burst as input to the classifier. Time points during reward delivery were excluded from the analysis to avoid the detection of the reward itself by the model. Negative samples (i.e. periods with no detected beta bursts) were randomly chosen time points of identical length (i.e., 50 frames), which did not overlap with the rewarded epochs. The ratio between positive and negative samples was kept at 1:1 for each session. The data was randomly separated (while keeping the ratio between positive and negative samples) into training and test sets for each training run (k-fold).

#### Support vector machine (SVM) classifier

To test for differences in behaviour during beta bursts epochs as compared to epochs without beta bursts, we employed a supervised linear classifier. To handle the large flow files (60 GB per 30 min session), we used an out-of-core incremental implementation of the support vector machines algorithm (sklearn^19^ 0.20.3).

The input data (samples X time points X frame-width X frame-length X motion-dimensions) were standard-scaled (mean subtracted and divided by the standard deviation, calculated pixel-wise on the training set) and linearized in a 1D array. Our model was implemented as Stochastic Gradient Descent Classifier (SGDClassifier^20^) with hinge loss and L2 regularization (alpha=0.0001). We evaluated the model accuracy as a mean of 3-fold splits of the data. Models with inputs of shuffled identity served as controls.

#### Model attention

Using a linear algorithm for classification allowed for projecting the model decision function back onto inputs and, thereby, for analysing the input subspace leading to the correct prediction. Model attention was defined as the distance of the input data point to the decision function. SVM fits a hyperplane (decision function) to the training data, which separates the classes with highest margin. Thus, for each data point, we calculated on which side of, and with which distance to the hyperplane the data point was located. Afterwards, we determined the time and space of the most important points, i.e. the points contributing the most to the correct classification. For representation purposes, attention was filtered with [0,2,3,3,0] Gaussian. Temporal attention was calculated as a sum of values over x, y, u and v dimensions per epoch and normalized to its maximum. Spatial attention was analysed for time points with maximal temporal attention within an epoch.

### Offline power and behavioural analysis

#### Group analysis

to demonstrate the flexibility of the method, we targeted two different ranges of beta oscillations: low-beta (15-20 Hz) in one rat and medium-high beta (20-25 Hz) in two rats (table S1). In order to investigate group effects, and due to the 1/f nature of LFP signals^21^, we calculated the normalized beta-power (Figs 2C, 2E and S2A) as follows; the power in each frequency in each session as computed online was averaged over the entire session (ignoring epochs of rejected artefacts). The session mean was then normalized to the mean of the first session, and the change of individual targeted beta frequencies was averaged and translated into percentages:

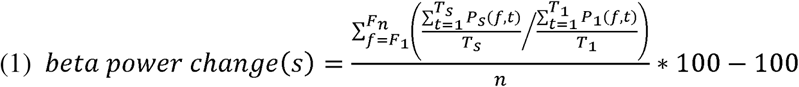

with s denoting the session, F = targeted frequencies, f = individual frequencies, n = 6 targeted frequencies, T = total number of time points in a session, t = individual time points (samples) and P= power. This value was averaged again over all rats to yield the group beta power change. Similar to power normalization, the number of rewards was normalized to the number of rewards in the first session (Figs 1E and S2B).

#### Within-subject analysis

to investigate the session-by-session and frequency-by-frequency effect in individual rats (figures S3 and S4), we used the mean (non-normalized) power of each targeted beta frequency in each session. For each rat, there was a session with a sudden, significant increase in beta-power (“aha-effect”). We used this session to group the days into two conditions (pre and post conditioning). In figure S4, the values are averaged over all “pre” sessions and all “post” sessions.

#### Comparison to other methods

We compared the proposed online filter-extrema method to 3 conventional time-frequency decomposition methods: wavelet, fast Fourier transform (FFT) and variance of filtered data (variation). For each method, two sets of parameters were used: “offline” as commonly used in offline analyses and “online” in which we applied computation-time constraints similar to our online methods (delay of 130 ms + half the period of each frequency). For each of the analyses, we divided the raw data into 100 “trials” (reward time-stamps ± 0.5 second) and padded with 1 second before and after the trial. The wavelet and FFT decompositions were computed using the fieldtrip toolbox^22^. For wavelet analyses, we used Morlet wavelets with width of 7 cycles (offline) or 3 cycles (online, corresponding to 130 ms + half the time period in 20 Hz), in steps of 1 ms. For FFT, we used Hanning time windows of 250 ms (offline) or 150 ms (online), in steps of 1 ms. The variation was computed by filtering the data (Matlab function “bandpass”, a Kaiser window FIR filter with stopband attenuation of 12 dB, passband of 1 Hz centred on the frequencies of interest and steepness of 0.75). Afterwards, the variance in each frequency was calculated over a window of 150 ms (offline) or half the time period of each frequency (online), sliding in 1 ms steps.

To compute the sum square error (SSE) metric, we normalized the power estimation of each method by the median of all trials. We applied the commonly used 7-cycle wavelet as the “gold-standard”, and subtracted each 2D matrix (time X frequency) of each trial from the wavelet values on this trial. The difference was squared and summed over time and frequency, as follows:

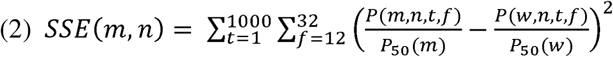

with m denoting the method, n- trial number (1 to 100), t- time point (ms), f- frequency (Hz), w- wavelet with 7 cycles width, P-power and P_50_- median power.

#### Statistical analysis

Statistical tests were performed in Matlab. For comparison of means of two groups (Fig. 2D), we used a two-tailed, two-sample t-test. To compare the SSE means of the different decomposition methods over trials (Fig. S1E), we used one-way analysis of variation (ANOVA). Two-way ANOVA was applied in figures 2C, S3 and S4. Whenever conducting multiple comparison post-hoc analyses, we used Bonferroni correction. To test linear regression, we computed Pearson’s correlations. The minimal number of animals needed for the study was determined using the resource equation approach^23^: Minimum n = ⌈10/(s–1)+1⌉, with n= 3, number of rats and s=9, number of sessions.

Code and example data sets are available on Github (https://github.com/arturoptophys).

---

**Figure S1:**
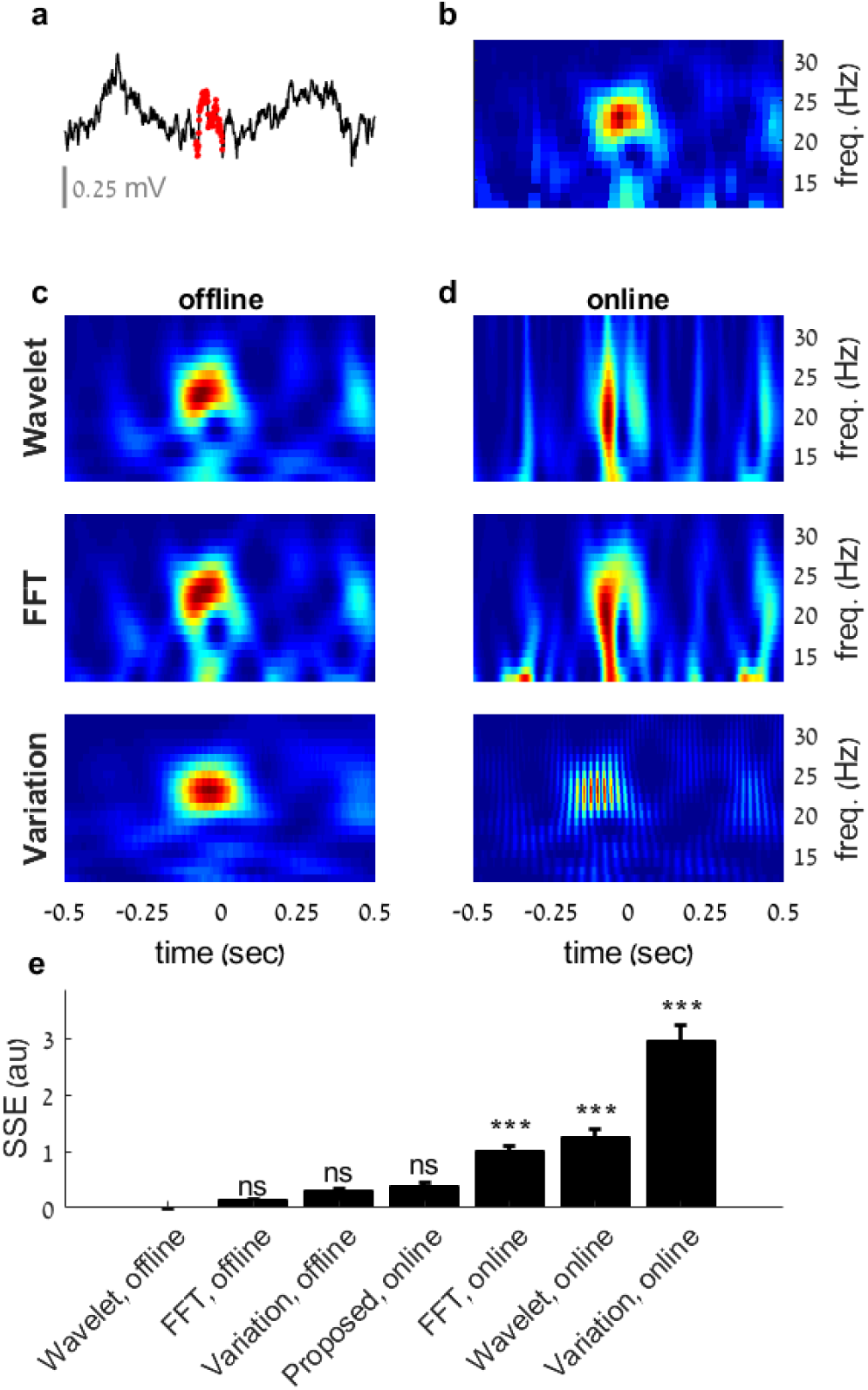
The filter-envelope method outperforms conventional methods in online beta-burst detection (related to figure 1). **a**. An LFP trace of a rewarded beta-burst (reward #98 in figure 2B) ± 0.5 sec. Time-points in which beta-power exceeded the 98th percentile threshold are marked in red. Reward was delivered at time = 0. **b**. Time-frequency decomposition of the trace in A using our online filter-extrema method. **c**. Time-frequency decomposition of the trace in A using three conventional offline analysis methods: convolution with Morlet wavelets with width of 7 periods (“Wavelet”, top), fast Fourier transform with sliding Hanning window sized 250 ms (“FFT”, middle) and the variance of the filtered data computed over windows of 150 ms (“Variation”, bottom). **d**. Applying the same time constraints as in the online filter-extrema method (delay of 130ms + half the period of each frequency) in the methods in C, resulted in worse resolution in the frequency dimension (Wavelet, top, and FFT, middle) or distortion in the time dimension (Variation, bottom). Wavelet and FFT were used with 3 periods and 150 ms accordingly to match the time delay for 20Hz. Variation was calculated over half the time period of each frequency. Note that for the wavelet, FFT and filter-extrema methods it is possible to extract the phase, while for the variation method, phase cannot be determined. **e**. The commonly used 7 periods wavelet was used as a standard, to which each method was compared to compute the sum of the square of the error (SSE) in 100 epochs of rewarded beta-bursts ± 0.5 sec. The proposed filter-extrema online method did not differ significantly from the offline methods, while all other online methods did (one-way ANOVA, F(6,693) = 74.006, p=2.87*10-71). Methods are sorted according to similarity to the offline wavelet method, and presented as mean ± SEM. ns-no significant difference in comparison to offline wavelet. ***-p<10-7, after Bonferroni correction for multiple comparisons.

**Figure S2:**
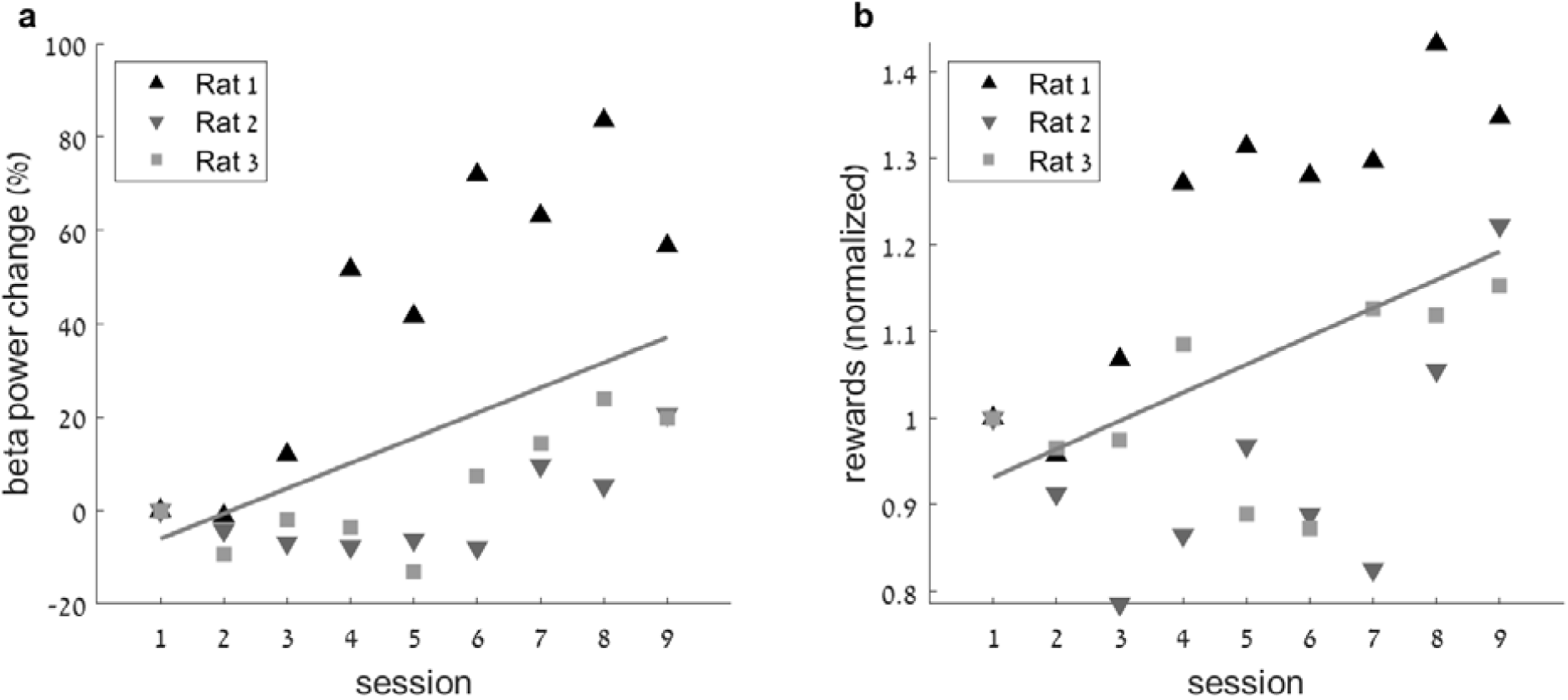
Beta power and rate increase by training (related to figure 2) Correlation between training session and **a**. beta power change (relative to day 1) and **b**. number of rewarded beta-bursts (relative to day 1). a. Pearson ρ=0.52, p=0.0059; b. Pearson ρ=0.49, p=0.0101.

**Figure S3:**
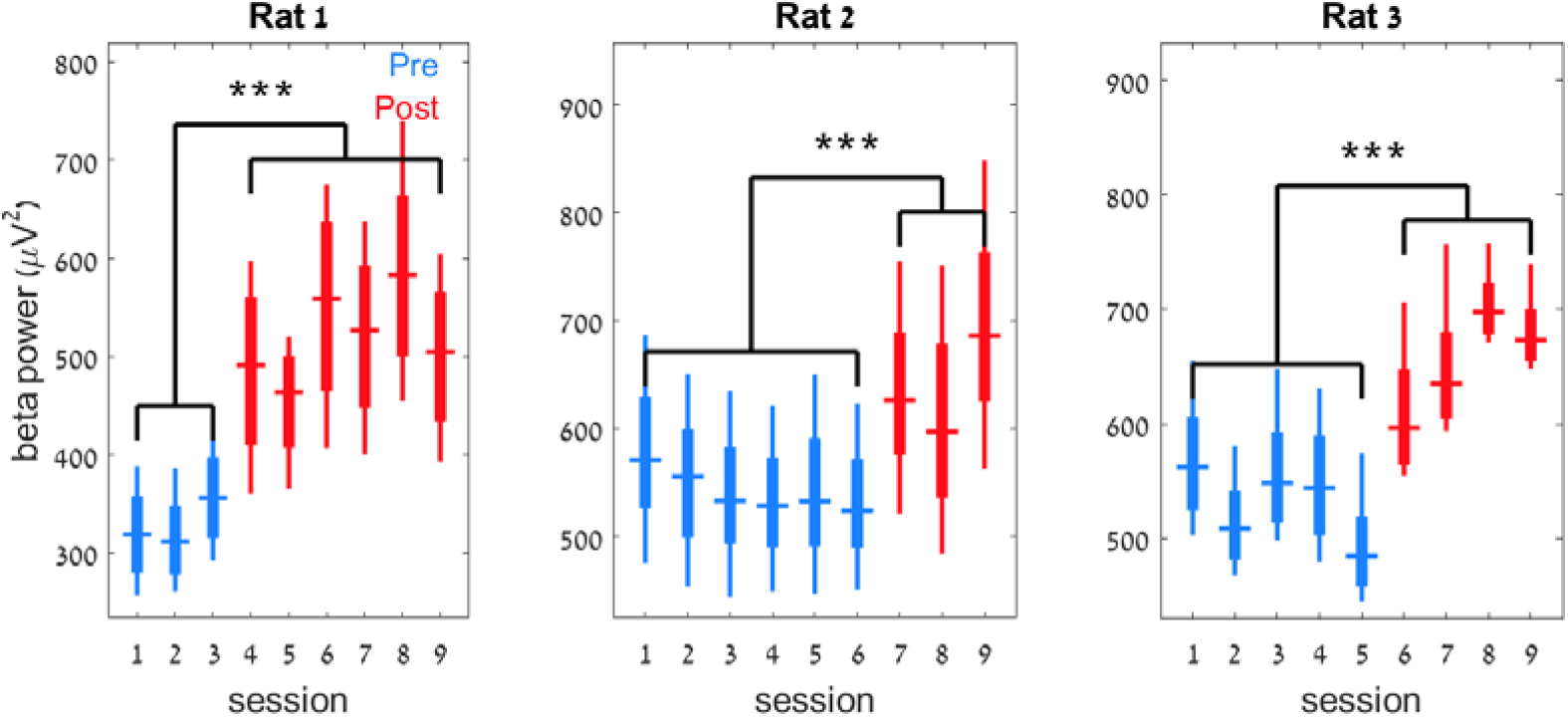
Rats exhibited an identifiable session of power increase – “aha-moment” (related to figure 2) The mean power of the targeted beta frequencies (20-25 Hz for rats 1 and 3, 15-20 Hz for rat 2) in each session. Note that for each rat there is a significant increase in power in a certain session that persists until the end of the experiment. Days before the increase are considered pre-conditioning (blue), and days after are considered post-conditioning (red). This definition implies also to figure 2D and figure S4. Two-way ANOVA (frequency and session), effect for session: rat1: F(5,8)=101.99, p=3.81*10-24, rat2: F(5,8)=85.73, p=10-22, rat3: F(5,8)=248.85, p=1.29*10-31, with Bonferroni correction for multiple comparisons between sessions. The presented elements are: centre line, median; box limits, upper and lower quartiles; whiskers, full distribution.***-p<5.19*10-6.

**Figure S4:**
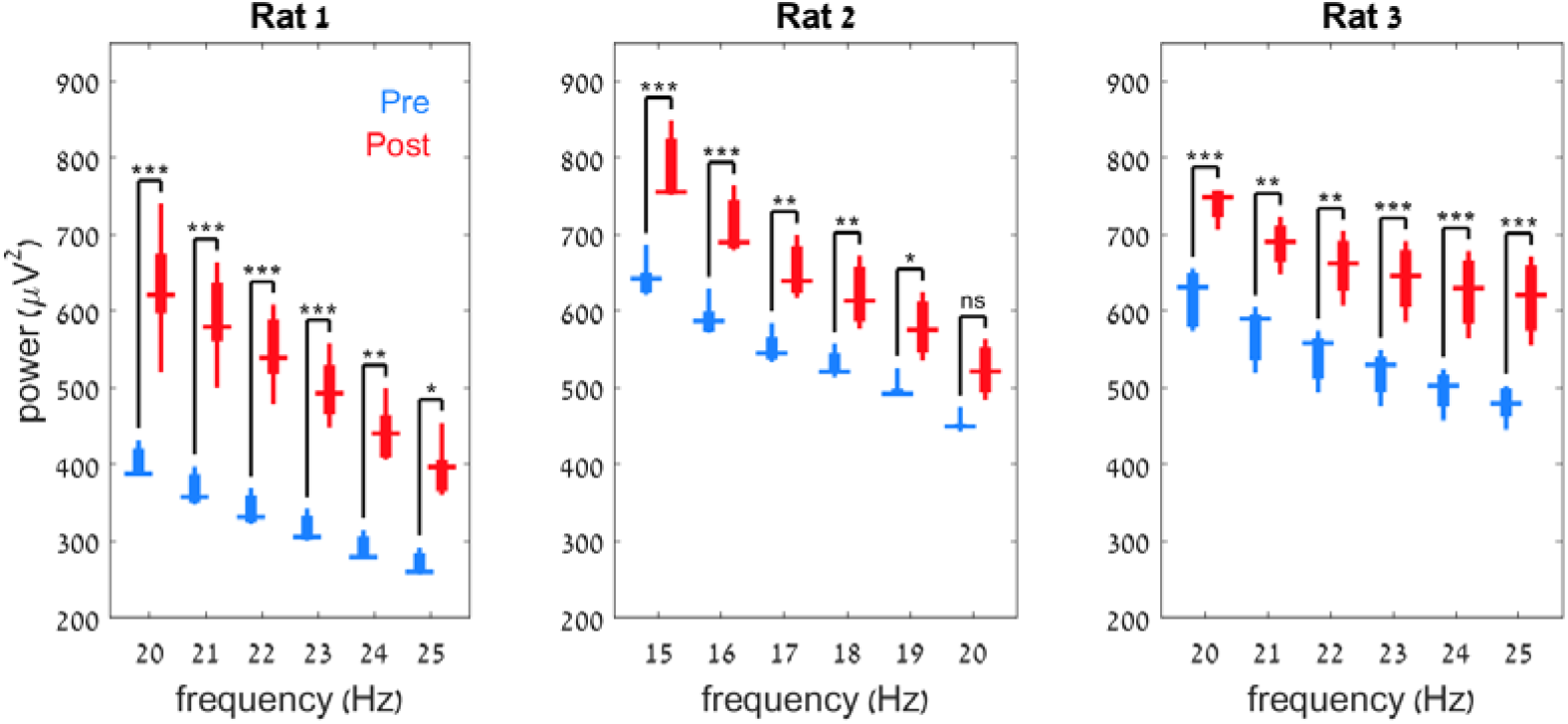
Increased power in individual frequencies by neurofeedback training (related to figure 2) The power of each of the targeted frequencies for sessions before (blue) vs. after conditioning (red). Two-way ANOVA (frequency and training state, pre or post conditioning), effect of training state: rat1: *F*_(5,1)_=191, p=5.08*10^−11^,rat2:*F*_(5,1)_=140,p=5.49*10^−15^, rat3: *F*_(5,1)_=152, p=1.510^−15^, with Bonferroni correction for multiple comparisons between the state in each frequency. The presented elements are: centre line, median; box limits, upper and lower quartiles; whiskers, full distribution.***-p<0.001, **-p<0.01, *-p<0.05, ns-not significant.

**Table S1:** details of the rats used in the study.

| **Rat number** | **1** | **2** | **3** |
| --- | --- | --- | --- |
| Rat designation number | 206 | 371 | 373 |
| Strain | Long Evans | Sprague Dawley | Sprague Dawley |
| Age at surgery (weeks) | 49 | 60 | 59 |
| Age at training initiation (weeks) | 94 | 65 | 65 |
| Weight at surgery (g.) | 370 | 360 | 322 |
| Weight at end of training (g.) | 380 | 350 | 310 |
| Session of "aha moment" | 4 | 7 | 6 |
| Target frequencies (Hz) | 20-25 | 15-20 | 20-25 |

**Table S2:** Sources of artefacts in electrophysiological recordings from freely moving rodents, and the measures to reduce their influence*.

| Artefact source | Artefact reduction measures |
| --- | --- |
| Alternating electric fields from the network supply (50/60 Hz “hum”) | Position the experimental setup in a grounded Faraday cage.<br>Uncouple the animal and recording pathway from the ground of the building (e.g. by optical interfaces). |
| Capacitance changes in moving electrical cables/ connections (“microphony”) | Digitize and amplify the signal at the headstage level.<br>Cement adaptors to avoid movement of connectors.<br>Keep the cable path clear of obstacles and tangling by an open-top cage and a commutator.<br>Reduce strain from the cable (in the “z” axis) by attaching soft springs to it.<br>Ensure clean and tight implant, with multiple reference points. |
| Electro-static discharges | To build the cage, use materials triboelectrically similar to the animal’s fur (e.g., glass). Avoid poly (methyl methacrylate), also known as Plexiglass or Perspex.<br>Wear a grounding wrist strap when handling the rats. |
| Alternating magnetic fields | Disconnect electrical transformers from sockets. |
| Switching devices (usually from the behavioural apparatus, e.g. lights and reward ports) | Keep high voltage devices out of the Faraday cage.<br>Use shielded and grounded cables. |
| Muscular activity, especially face muscles | To reduce epochs of teeth crunching, maintain the subject motivated and satisfied (e.g. by adjusting the reward size).<br>Detect epochs of facial muscles activity/ touching the headstage and remove them from analysis. |
| Direct-current discharges from touching the ground | Keep the rat and recording system floating above ground (by isolating materials e.g. glass and wood).<br>Keep large conductive materials out of the cage. |

\* see also: https://neuralynx.com/news/techtips

